# Mutant KRAS promotes NF-κB driven CCL20 chemokine expression in pancreatic ductal adenocarcinoma

**DOI:** 10.64898/2026.04.13.717530

**Authors:** Donovan Drouillard, Marissa Davies, Donna McAllister, Michael B. Dwinell

## Abstract

The chemokine CCL20 is implicated in inflammation and cancer but has proven challenging to target therapeutically. In this study, we precisely define what cells produce CCL20 in pancreatic inflammation and cancer. Through analysis of single cell RNA data, mutation and copy number signatures, gene methylation, and *in vitro* studies, we show that CCL20 and other NF-κB driven chemokine production is largely dependent on oncogenic KRAS in the malignant pancreas. Blockade of CCL20-CCR6 signaling *in vivo* using a novel partial agonist inhibitor, CCL20LD, increased recruitment of antigen presenting cells without significantly impinging tumor growth. Lastly, resistance to pan-RAS or allele-specific KRAS inhibitors decreased CCL20-dependent immune recruitment in culture. These results suggest that oncogenic KRAS activates NF-κB signaling in human pancreas cancer, resulting in pharmacologically reversible changes to chemokine production that may participate in immune suppression or immune evasion within the pancreas cancer microenvironment.

## INTRODUCTION

Chemokines are a subset of cytokines that direct leukocyte recruitment during homeostasis and inflammation. Accordingly, homeostatic chemokine ligands are expressed in a healthy state to maintain immune surveillance while inflammatory chemokines are expressed largely following inflammatory cues such as bacteria or viral infection and/or tissue damage^1^. CCL20, the only cognate ligand for the chemokine receptor CCR6, falls in between these two broad classifications. In a healthy state CCL20 is constitutively expressed by the follicle associated epithelium in Peyer’s patches and isolated lymphoid follicles along the GI tract, where it is believed to facilitate homeostasis between the gut microbiome and the host^2,3^. However, during infectious or sterile inflammation, CCL20 expression is upregulated across a wide range of cells including epithelial cells, keratinocytes, Th17 cells, and macrophages^4^. Dysregulation of constitutive CCL20 expression during inflammation is thought to contribute to the pathophysiology of many autoimmune diseases including psoriasis, inflammatory bowel disease (IBD), rheumatoid arthritis, and multiple sclerosis^5–7^. CCL20 has been linked indirectly to damage in the associated epithelial or neuronal tissue in those disease states through recruitment of CCR6+ Th17 cells and subsets of regulatory T cells (Tregs), B cells, dendritic cells, and macrophages that release cytokines, granzymes, and other effector molecules that invoke dysfunction in the epithelia or nearby neurons^8,9^. Paradoxically, CCR6-expressing Th17 cells produce proinflammatory cytokines which cumulatively perpetuate fibrotic inflammation in the diseased gut while peripheral regulatory T cells (Treg) maintain self-tolerance and prevent overactive fibroinflammatory reactions in autoimmunity. While immune tolerance reflects the homeostatic balance in Th17 and Treg cell populations, autoimmune reactions in colitis reflect an imbalance where proinflammatory Th17 cell populations overwhelm protective Treg levels in the mucosa. An unresolved question is how CCR6+ Th17 and CCR6+ Treg populations become unbalanced as the concentration of CCL20 shifts from a healthy to a diseased mucosa.

The formation of the fibrotic immunosuppressive tumor microenvironment (TME) that occurs during pancreatic ductal adenocarcinoma (PDAC) development is a major contributor to the poor prognosis of the disease and its poor response to immunotherapy^10^. In PDAC, only *BRCA1/2*-mutant and mismatch repair-deficient patients, which combined account for less than 10% of diagnosed PDAC patient tumors^11,12^, have shown strong responsiveness to immune checkpoint inhibitor immunotherapy^13,14^. Prevention or disruption of the immunosuppressive TME may provide an opportunity to sensitize more patients to immunotherapy. One likely method of disrupting the TME is through modulation of chemokine signaling, as seen in clinical trials using either a dual CXCR1/2 inhibitor (NCT04477343) or CXCR4 inhibitor (NCT04177810) in patients with metastatic PDAC. To date, several clinical trials targeting the chemokine CXCR4 have failed to show any benefit, likely reflecting the near ubiquitous expression profile of the receptor^15^. While no data have yet to be published on the clinical efficacy of CXCR1/2 inhibition in PDAC, preclinical data show that CXCR1/2 inhibition increased T cell infiltration, decreased myeloid derived suppressor cell infiltration, and synergized with anti-PD-L1 blockade^16^. However, while production of CXCR1/2 ligands is elevated in some PDAC patient tissues, the receptor for these chemokines is restricted to CXCR1 and CXCR2 expressing granulocytic immune cells. Thus, CXCR1/2 immunotherapy may not fully account for the other disparate and abundant immune cell types within the TME that enforce immune suppression such as Tregs. Further, differences in tumor subtype, oncogene dosing, tumor mutation burden, and other genetic and non-genetic factors have all been shown influence the composition of the TME^17^. Thus, there remains a need to more fully examine non-CXCR1/2 ligands and better understand the mechanisms that drive aberrant chemokine expression to identify precision immunotherapies for PDAC. Collaborative efforts in our group have uncovered crucial molecular mechanisms of chemokine biology that may help explain discrepancies in Th17 and Treg migration into diseased tissues^6,8,18–20^.

CCL20 upregulation has been variably linked in response to malignant EGFR overactivation^21^, promoter hypomethylation within the CCL20 promoter region^22^, and 5-fluorouracil or taxane chemotherapy use^23,24^ in multiple different cancer types and preclinical models. Most studies postulate that the CCL20-CCR6 signaling axis is detrimental in cancer^21,22,25,26^, without a consensus mechanism or cellular target for the putative tumor-promoting phenotype. Production of CCL20 has been shown to be dysregulated in PDAC, although the cause and effects of aberrant CCL20 in malignancy remains little understood^27^. Implicated mechanisms whereby altered CCL20 may promote tumor progression include recruitment of endothelial cells during tumor neovascularization^21^, tumor infiltration of Tregs^23^ and/or Th17 T cells^22^, or direct growth or migration effects on tumor cells^26^. Further uncertainty stems from the ambiguous role of Th17 cells in cancer. Th17 cells have been varyingly shown to directly influence cancer initiation^28^, provide either anti- or pro-tumorigenic effects in the tumor microenvironment^29,30^, or play a direct role in tumor elimination through induction of tumor-reactive antibodies in B cells^31^. Given these persistent gaps, there remains a critical need to define when and where CCL20 is upregulated in PDAC, which cells CCL20 recruits within the heterogenous tumor microenvironment, and how these alterations ultimately impact tumor progression and immune control.

In this study, we used an unbiased approach to identify the CCL20 producing cells in healthy, inflamed, precancerous, and malignant pancreas and colon through mining of single cell RNA datasets and investigated the mechanisms regulating CCL20’s expression. Using single cell RNA and The Cancer Genome Atlas (TCGA) data, we precisely identified the cancer subtypes, mutational signatures, and epigenetic changes associated with increased CCL20 expression. Comprehensive screening approaches of patient samples was confirmed in PDAC cell lines and demonstrated that tumor cells are the predominant producers of CCL20. Further, tumor cell CCL20 expression was dependent on mutant KRAS. Lastly, we used a novel first in-class inhibitor of CCR6 that we have previously shown to ameliorate Th17 T cell recruitment in preclinical models of dermal autoimmune inflammatory diseases to inhibit the tumor infiltration of antigen presenting cells within PDAC tumors *in vivo*. These data indicate that blocking CCR6-target cells promotes the infiltration of antigen presenting myeloid subsets into murine PDAC tumors with oncogenic KRAS mutations.

## RESULTS

### CCL20 upregulation in gastrointestinal malignancies

Given CCL20’s known restriction to mucosal, relative to epidermal, tissues, we interrogated CCL20 expression at a pan-cancer level using The Cancer Genome Atlas (TCGA)^32^ and Genotype-Tissue Expression (GTEx)^33^ project data. CCL20 is significantly upregulated in all gastrointestinal (GI) malignancies compared to their respective healthy tissues (Fig. 1A). Other solid and hematological malignancies had little to no CCL20 expression, with some non-GI, mucosal malignancies upregulating CCL20. To validate these findings, we analyzed single cell RNA (scRNA) datasets of human PDAC^34^ and colorectal cancer (CRC)^35^. In both PDAC and CRC, CCL20 was expressed by the myeloid and epithelial populations (Fig. 1B, 1C). Further analysis of the CCL20+ myeloid cells in both PDAC and CRC showed expression limited to a subset of IL-1β+ macrophages (Supplementary Data 1, Supplementary Data 2). Inflammation upregulates CCL20 expression and is a risk factor for both PDAC and CRC^36^, so we examined human pancreatitis^37,38^ and colitis^39^ scRNA and single-nuclear RNA (snRNA) datasets to investigate if CCL20 was equally upregulated in the inflamed, non-malignant state. In the pancreatitis datasets, we found only 5% of a subset of mucin-producing ductal cells expressed CCL20 suggesting CCL20 upregulation by pancreatic epithelium requires malignant transformation (Fig. 1D). In contrast to PDAC tumors, increased CCL20 expression in pancreatitis patients was localized largely to CD4+ T cells and M2-like macrophages, while IL-23A and IL-17A were rarer and expression was limited to a subset of CD4 T cells (Fig. 1E). Analysis of the colitis dataset showed comparable myeloid upregulation of CCL20 (Fig. 1F), with colon epithelial cells in ulcerative colitis and Crohn’s disease expressing a higher baseline expression of CCL20 compared to ductal cells in pancreatitis. These findings are in line with previous reports of high CCL20 production by intestinal epithelium when given a variety of inflammatory stimuli and are congruent with the presence of a large microbiome in the gut relative to the pancreas^20^.

**Figure 1.**
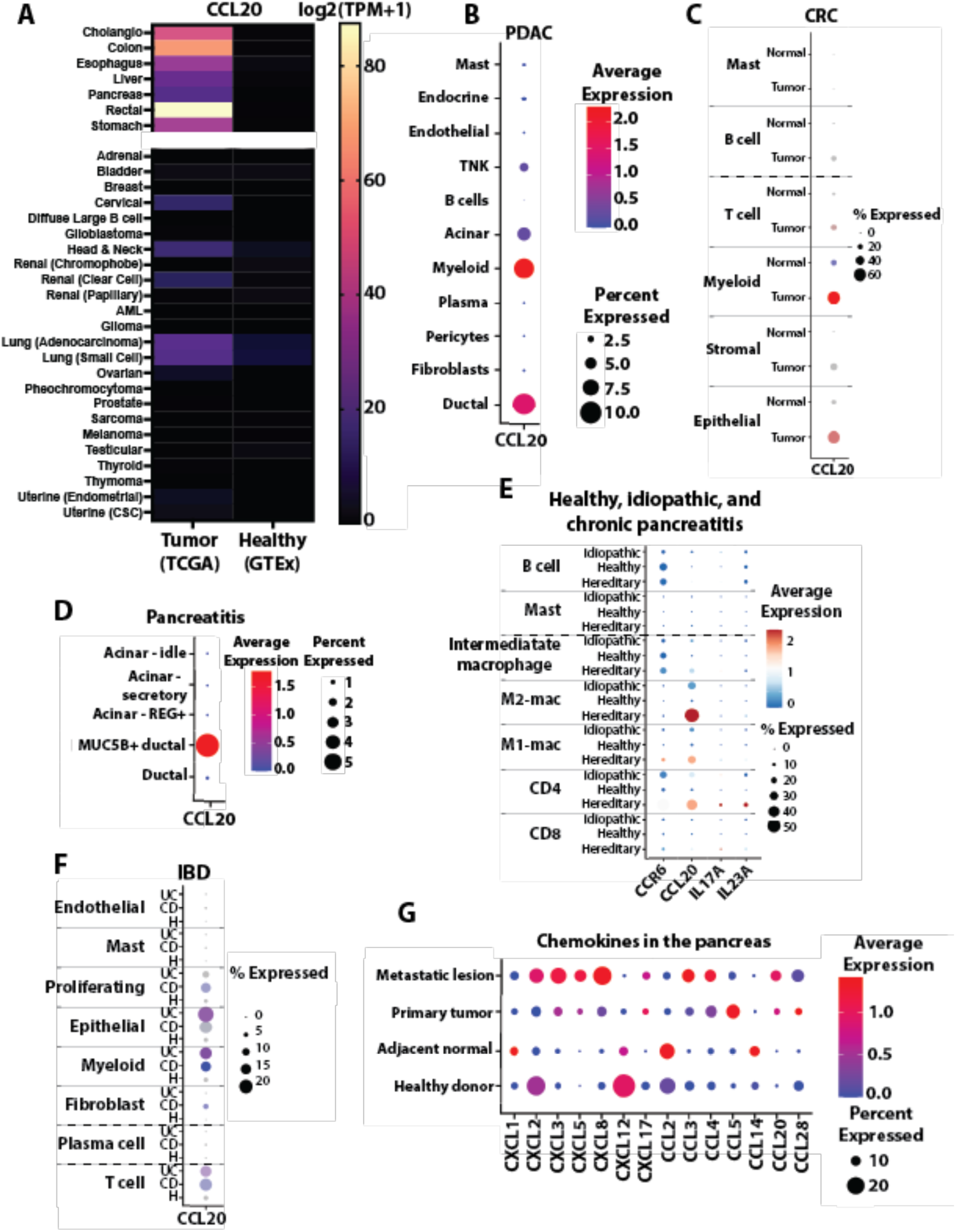
CCL20 is upregulated in human PDAC and chronic pancreatitis. **A** Bulk sequencing data from TCGA (left) and GTEx (right) showing CCL20 expression. Gastrointestinal tissues prominently express CCL20 while other epithelial tissues expressed little if any CCL20. **B** Expression of CCL20 by the different cell types found in primary tumors from a PDAC scRNA atlas^34^. **C** CCL20 expression from a scRNA dataset healthy human colon (H) and colorectal cancer^35^. **D** Expression of CCL20 by different cells found in a single nucleus RNA sequencing dataset of chronic pancreatitis biopsies^38^. **E** CCR6, CCL20, IL-17A, and IL-23A in a scRNA datasets of CD45+ cells from human chronic pancreatitis or healthy patients^37^. Pancreatitis was classified as hereditary if there were mutations present in *PRSS1* gene. **F** Expression of CCL20 in a scRNA dataset of human ulcerative colitis (UC) and Crohn’s disease (CD)^39^. **G** Differentially expressed chemokines found using MAST^103^ from a scRNA atlas of PDAC^34^.

To further investigate the distinct chemokine profile of PDAC that mediates formation of the immunosuppressive TME, we investigated differentially expressed chemokines in healthy, adjacent normal, primary tumor, and metastatic lesions of the pancreas (Fig. 1G)^34^. Homeostatically expressed chemokines in the healthy and adjacent normal pancreas were CXCL12, CCL2, and CCL14. Of these, CXCL12 is a hallmark homeostatic chemokine, while CCL2 and CCL14 are two monocyte-specific chemoattractants that may play role in trafficking of tissue resident macrophages to the exocrine pancreatic mucosa. CXCL1, a proinflammatory chemokine, as well as CCL2 and CCL14, were upregulated in the adjacent normal pancreas. While not statistically significant the reduction in CXCL12 from healthy to adjacent normal pancreas, which further extinguishes in primary and metastatic PDAC, aligns with our prior work demonstrating its epigenetic silencing in malignant cancers^40,41^. These data are consistent with the limited immune cell infiltration within healthy pancreas. The transient increase in the neutrophil chemoattractant CXCL1 is suggestive of playing a role early in malignant transformation. Differentially expressed chemokines in both metastatic and primary tumors included the neutrophil chemoattractants CXCL3 and CXCL5, as well as CCL20. Intriguingly, metastatic lesions had a heterogenous mix of chemokine expression that included neutrophil chemokines CXCL2 and CXCL8, the NK targeting chemokine CXCL16, as well as CCL3 and CCL4 chemokines that play roles in monocyte recruitment. The CXCR1 and CXCR2 ligands, CXCL1-3 and CXCL5-8, have been well studied for their role in the PDAC TME. Additional chemokines identified in primary tumors were CXCL14^42^, CXCL17^43^, CCL5^44^, and, consistent with our prior report, CCL28^45^. While elevated expression of CCL20 has been observed, its functional role in PDAC progression and immune remodeling remains little understood, with paradoxical roles either promoting or inhibiting tumor progression noted. CCR6-expressing Th17 cells are typically associated with CCL20, however the Th17 cytokines IL-17A, IL-17F, IL-21, and IL-22 were not differentially expressed between healthy and malignant pancreas suggesting non-Th17 CCR6+ cells were being recruited to PDAC tumors (Supplementary Data 3).

### CCL20 is upregulated in model systems of PDAC

To confirm that CCL20 is upregulated in PDAC cell lines, we screened both human and mouse cell lines for CCL20 by PCR. Expression was detected in nearly all PDAC cell lines, including two patient-derived PDAC cell lines (Fig. 2A). The MiaPaCa2 PDAC cell line was CCL20 negative, matching previous reports which found the cell line to have mutant USP15, a deubiquitinase that regulates NF-κB signaling which may be necessary for CCL20 transcript expression^46^. Testing of mouse KPC (LSL-*Kras*^G12D/+^; LSL-*Trp53*^R172H.+^; *Pdx1*-Cre) PDAC cell lines showed consistent expression of CCL20 (Fig 2B). In contrast to the ligand, transcript expression of CCR6, the receptor for CCL20, was faintly if at all detected in PDAC cell lines. Immunofluorescence staining of healthy human pancreas and human PDAC showed significant CCL20+ staining in PDAC samples but little to no CCL20 staining in healthy pancreas (Fig. 2C) and as indicated in representative immunofluorescence images (Fig. 2D, 2E). Together, these data underscore that the ligand is aberrantly expressed by epithelial cells in the fibro-inflamed and malignant pancreas of humans and the relevant preclinical models and confirm the transformed epithelium as a predominant source of the elevated CCL20 production. The lack of CCR6 expression by the malignant tumor cells indicate that the tumors are not the most likely target for chemokines in PDAC.

**Figure 2.**
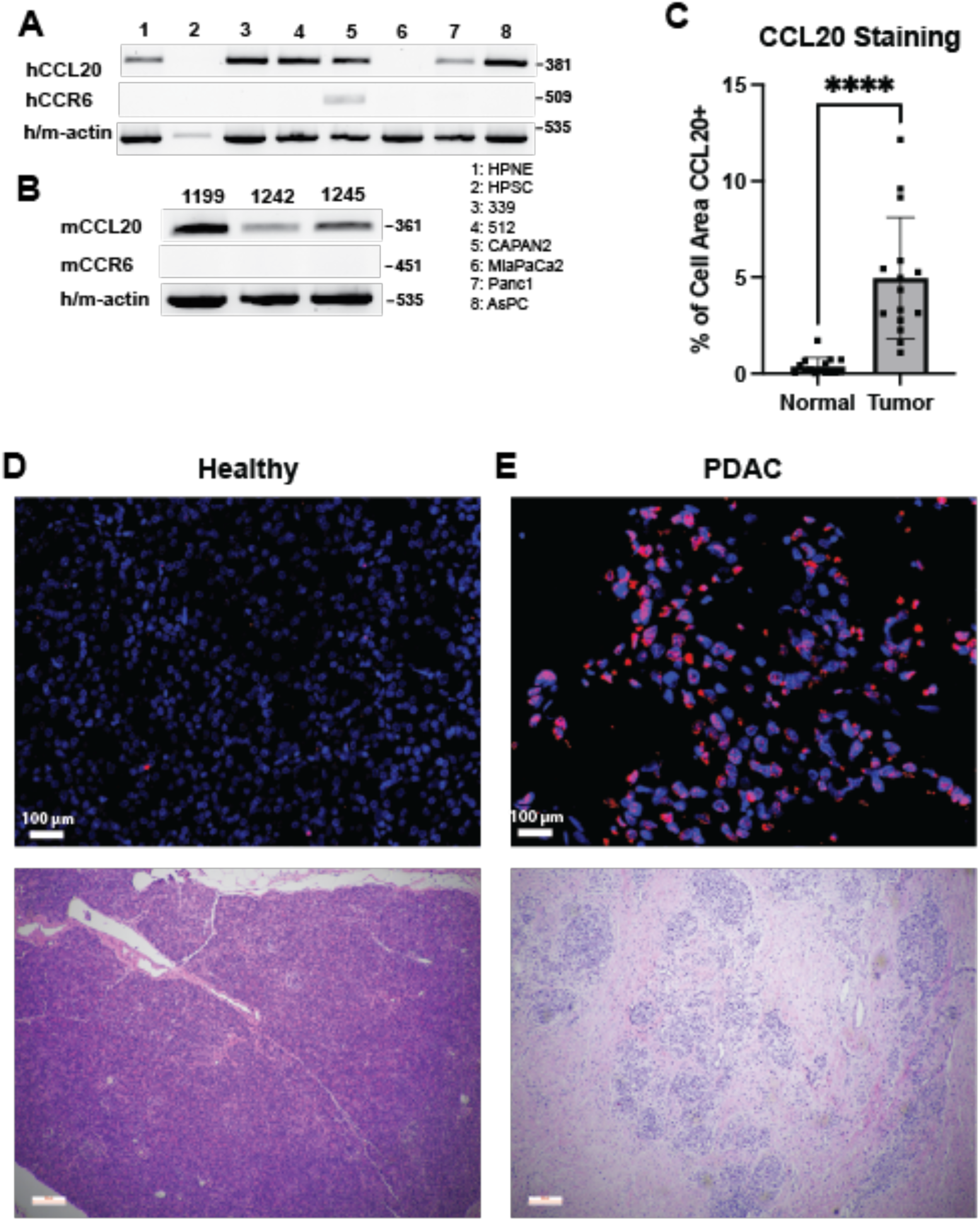
CCL20 expression in PDAC cell lines and human healthy pancreas tumor sections. **A** Representative PCR analyses of CCL20 and CCR6 transcript expression across multiple human and **B** mouse PDAC cell lines. Data representative of 3 individual analyses. **C** Quantification of the CCL20 staining images. 3-7 fields of view were quantified per patient, with five patients per disease state**. D** Representative immunostaining for CCL20 (red) or DAPI-stained cell nuclei (blue) (top) with parallel sections stained with H&E (bottom) in both healthy human pancreas or **E** human PDAC samples. * denotes *p* ≤ 0.05, Student’s unpaired t-test. Size bar = 100 μm.

### Tumor subtype specific analysis of CCL20 expression

Classification of PDAC and CRC tumors cells into different subtypes has shown meaningful impact on prognosis and is beginning to impact the therapies given (NCT04683315)^47–49^. To further explore the mechanism behind malignant upregulation of CCL20 specific to PDAC but not CRC, we used the previously mentioned scRNA datasets to explore correlated genes and cell programs. We discovered that in PDAC, CCL20 expression was correlated with the basal, and not the classical or intermediate, forms of PDAC using previously established sets of transcripts (hereafter referred to as gene sets) to distinguish between subtypes (Fig. 3A)^50^. The basal subtype of PDAC frequently undergoes epithelial to mesenchymal transition (EMT) and in line with these findings, we also observed a positive correlation between EMT gene sets and CCL20 expression^51^. Other gene sets positively correlated with CCL20 in PDAC included receptor tyrosine kinase (RTK) and EGFR signaling, cellular programs typically upregulated in cancer. However, other cellular programs upregulated with cancer such as chromothripsis, extracellular vesicle mediated signaling, and the senescence-associated secretory phenotype (SASP) had negative correlations with CCL20. The same gene sets were examined for correlation with CCL20 expression in the CRC scRNA dataset to see if there was a pan-cancer gene set responsible for CCL20 expression. We demonstrated that the iCMS2 subtype of CRC, and not the iCMS3 subtype, was primarily associated with CCL20 expression (Fig. 3B)^48^. In contrast with the basal PDAC subtype and CRC iCMS3 subtype, the iCMS2 subtype of CRC is well-differentiated, as validated by the negative correlation between CCL20 and EMT in CRC^52^. Additionally, the iCMS2 CRC subtype is associated with a favorable prognosis^48^ while the basal PDAC subtype is associated with a poor prognosis further highlighting the dichotomy between PDAC and CRC CCL20 expression^47^. We observed a similar trend with CCL20, which was a negative prognostic marker in PDAC and positive prognostic marker in CRC (Fig. S1A, S1B). The iCMS2 subtype expression of CCL20 may be explained by its predominance in the left colon, which has been found to harbor more pathogenic bacteria when tumors are present compared to the right colon^53^. Thus, CCL20 elevation was correlated with a basal, more de-differentiated subtype in PDAC compared to a well-differentiated form of CRC whose expression of CCL20 may be dependent on external cues. Our observed differences between PDAC and CRC CCL20 production may explain the lack of a consensus mechanism for CCL20’s tumor promoting attributes and highlight the need for more precise study of CCL20 pathophysiology in cancer.

**Figure 3.**
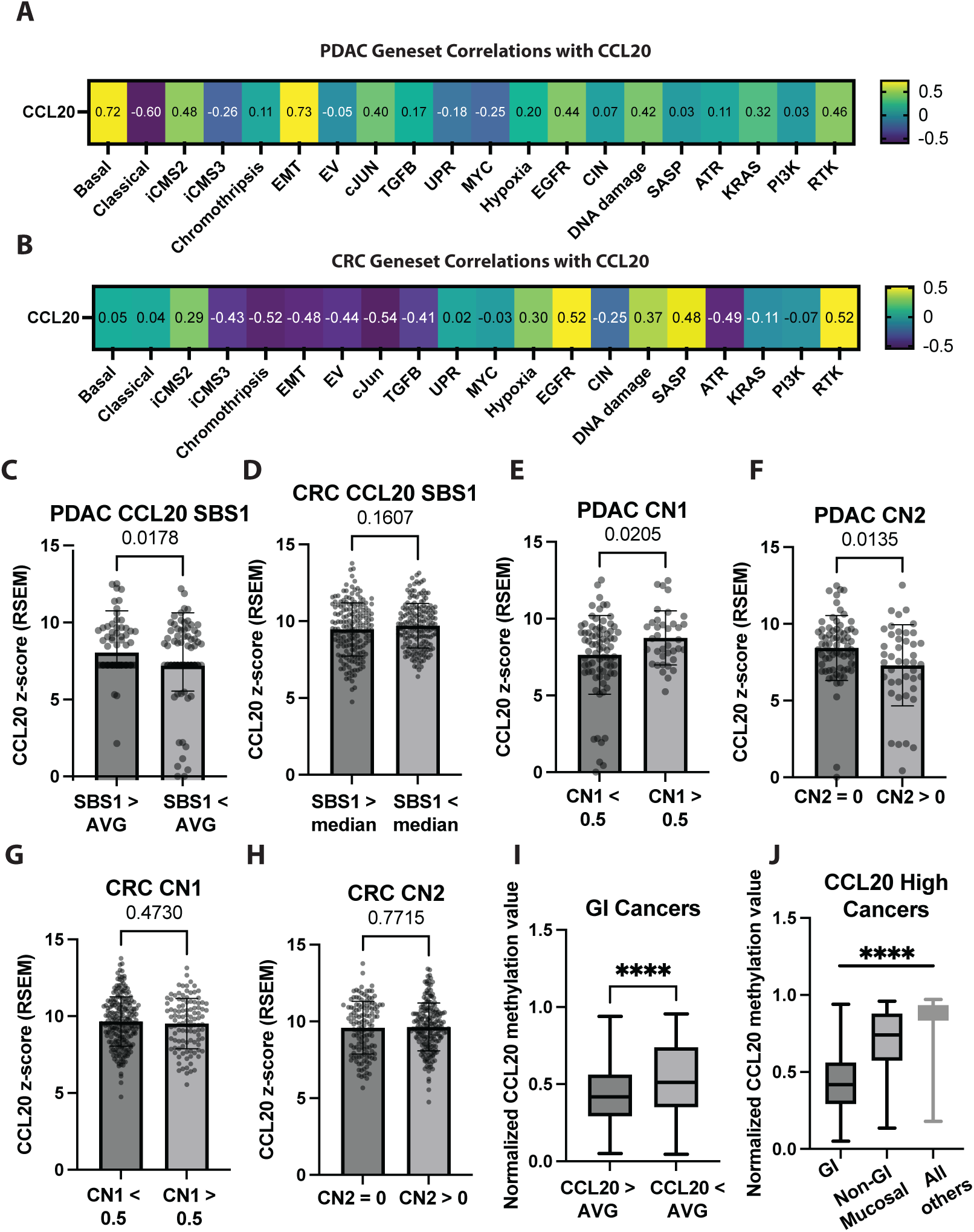
Gene sets, mutation signatures, and copy number signatures associated with CCL20. **A** Gene sets listed in Supplementary Data 7 were correlated with CCL20 expression in ductal cells from a scRNA dataset of human PDAC^110^ and **B** colorectal cancer^35^. Values shown are Pearson’s correlation coefficients, all p values = 0. **C** TCGA PDAC patients and **D** colorectal cancer patients sorted by SBS1 (replicative age) signatures, with comparisons of their normalized CCL20 expression. **E** TCGA PDAC patients sorted by their CN1 (diploid) and **F** CN2 (tetraploid) signatures using data from previously reported^58^. Similar analysis for colorectal cancer patients can be seen in **G** and **H**. **I** Normalized CCL20 methylation values (probe cg21643045) from all GI cancers (cancer types in the top row of Fig. 1E) sorted by CCL20 expression. **J** CCL20 high patients (relative to patients of same cancer type) with their CCL20 normalized methylation values plotted. Non-GI mucosal cancers are bladder, cervical, head and neck, both lung types, and both uterine tumor types. Panel C-I Student’s unpaired *t*-test was used with *p* values plotted. Panel J one-way ANOVA with Tukey’s test for multiple comparisons, * denotes *p* ≤ 0.05.

### Mutation and copy number signatures associated with CCL20

Having confirmed subtype specific expression of CCL20 in both PDAC and CRC, we next investigated mutation signatures^54^ to see if there was a shared pattern of DNA damage associated with CCL20 expression. For these studies, we analyzed mutation signatures correlated with elevated CCL20 from TCGA using the publicly available somatic mutation annotation format (MAF) files. Single base substitution (SBS) signatures detected in >10% of PDAC TCGA samples included SBS1 (deamination of 5-methylcytosine), SBS5 (unknown, clock-like), SBS10b (polymerase epsilon exonuclease domain mutations), SBS15 (defective DNA mismatch repair), and SBS87 (thiopurine chemotherapy treatment) (Supplementary Data 4). The same SBS signatures, plus SBS6 (defective DNA mismatch repair), were detected in >10% of CRC TCGA samples (Supplementary Data 5). Confirming that bulk TCGA gene expression correctly correlates with the SBS signatures, we found increased expression of APEX1 (apurinic/apyrimidinic endodeoxyribonuclease 1) and LIG1 (DNA ligase 1) in SBS1 high samples in both PDAC and CRC (Fig. S2A). APEX1 and LIG1 are important in the base excision repair pathways responsible for fixing 5-methylcytosine deaminations^55^, thus proving that expected increases in bulk gene expression could be observed based on separation by mutational signatures. Only SBS1 was correlated with increased CCL20 expression in PDAC, while no SBS signatures, including SBS1, were correlated with CCL20 expression in CRC (Fig. 3C, 3D). SBS1 and SBS5 have both been termed “clock-like” due to the observation of their linear accumulation with age^56^. However, recent data suggest that the SBS1 signature is driven by replicative age while the SBS5 signature accumulates independently of cell divisions and is truly clock-like^57^. In line with these data, we measured within the PDAC TCGA dataset that the SBS1 high group had a higher frequency of KRAS mutations but no significant correlation with age, while the SBS5 high group trended toward significance with age, with only a modest difference in KRAS mutation frequency (Fig. S2B). No differences in KRAS mutation frequency were observed between different SBS signatures in the CRC dataset (Fig. S2C).

Accurate copy number (CN) signature analysis requires whole genome sequencing data, which was not publicly available through the TCGA data on the GDC data portal. However, Steele et al. recently generated CN signatures for all TCGA samples^58^. Using their CN signature data, the CN signatures detected in >10% of PDAC samples included CN1 (diploidy), CN2 (tetraploidy), and CN9 (focal loss of heterozygosity – diploid and chromosomal instability). CRC had the same CN signatures detected in >10% of samples, in addition to CN3 (octoploidy) and CN17 (tandem duplication and homologous repair deficiency). Only CN1 and CN2 were positively and negatively associated with CCL20 expression in PDAC, respectively (Fig. 3E, F). No significant associations were observed in CRC (Fig. 3G, 3H). This suggests two possibilities for PDAC. One explanation is that whole or focal genome duplications are negatively associated with CCL20 expression in PDAC. Another explanation is that maintenance of a diploid genome is positively associated with CCL20 expression in PDAC. Due to the trending positive association with CN9, focal loss of heterozygosity, it is likely that a diploid genome with focal losses would result in the highest CCL20 expression in PDAC. However, CCL20 did not reach significance when correcting for multiple comparisons for any copy number signature. Together, these data suggest that an absence of a unifying pan-cancer genome alteration associated with malignant transformation influencing CCL20 aberrant expression in cancer.

### Pan-cancer CCL20 gene methylation patterns

Given the limited genetic changes correlated with CCL20’s increased expression we next asked if epigenetic changes may be another mechanism of differential CCL20 expression in cancer^22^. To study this mechanism, we looked at methylation of the CCL20 gene across all mucosal cancers using cBioPortal^59^. We found decreased CCL20 methylation in GI TCGA samples with high CCL20 expression compared to GI TCGA samples with low CCL20 expression (Fig. 3I). Across all CCL20-high cancers, we found TCGA GI tumors had the lowest CCL20 gene methylation, followed by TCGA non-GI, but mucosal tumor samples; and non-GI, non-mucosal, TCGA tumor samples having the highest normalized CCL20 gene methylation (Fig. 3J). These data suggest that changes in CCL20 transcript expression may be influenced through cancer-mediated epigenetic changes rather than genomic alterations.

To further explore the possibility of epigenetic regulation of CCL20 in cancer, we used the HCT116 CRC cell line which possesses a KRAS G13D mutation. We compared CCL20 transcript expression in HCT116 wild-type cells with HCT116 cells in which DNA methyltransferase (DNMT) 1, DNMT3b, or both had been deleted^41^. The HCT116 cell line had low constitutive CCL20 expression and protein (Fig. S3A, S3B), consistent with The Human Protein Atlas data for CRC (Fig. S3C). However, these DNMT knockout cells retained the ability to respond to proinflammatory cytokines, with either TNF or IL-1 stimulating CCL20 protein production. Knockouts of DNMT1, DNMT3b, or both resulted in minor decreases in basal expression and no changes in protein levels of CCL20. This contrasts the data seen in Figures 3I and 3J, suggesting that decreases in DNA methylation in the presence of the KRAS G13D mutation are not enough to elevate CCL20 expression. Interestingly, CXCL2 expression did not follow the same patterns. Sole knockout of DNMT1 or DNMT3b was sufficient to lower basal CXCL2 expression in the HCT116 cell line, while the dual DNMT1+3b double knockout resulted in a minor increase in CXCL2 expression (Fig. S3D). Together, these data indicate that epigenetic modification of the CCL20 gene locus may negatively influence expression, with other factors likely play larger roles in altering its transcriptional expression.

### Transcriptional programs regulating CCL20

Due to the lack of consistent genetic and epigenetic changes we found associated with CCL20, we next examined transcription factors that could be driving CCL20 expression. To do so, we computationally inferred transcription factors^60^ that were differentially active in CCL20+ compared to CCL20 negative PDAC tumors cells from a scRNA dataset^61^. The results suggested that components of NF-κB signaling were driving CCL20 gene expression in PDAC tumor cells (Fig. 4A). In line with our data in Figure 3A, CCL20 negative PDAC tumor cells had a higher PDX1 score, a transcription factor important for pancreas differentiation^62^ suggesting that CCL20+ PDAC tumor cells are less differentiated. The NF-κB transcription factor family was also implicated in CCL20+ CRC tumor cells (Fig. 4B). To confirm these results in a cell agnostic manner, we used ContraV3^63^, a tool to predict transcription factor binding which further supported our results (Fig. S4A). Lastly, other NF-κB regulated chemokines, such as CXCL2^64^ and CXCL3^65^ were upregulated in both the CCL20+ PDAC and CRC tumor cells compared to their CCL20 negative counterparts (Fig. S4B, S4C). Altogether, these data suggest that while the initial provocation to upregulate CCL20 may be different between PDAC and CRC, the resulting signaling pathways both converge on NF-κB to upregulate CCL20.

**Figure 4.**
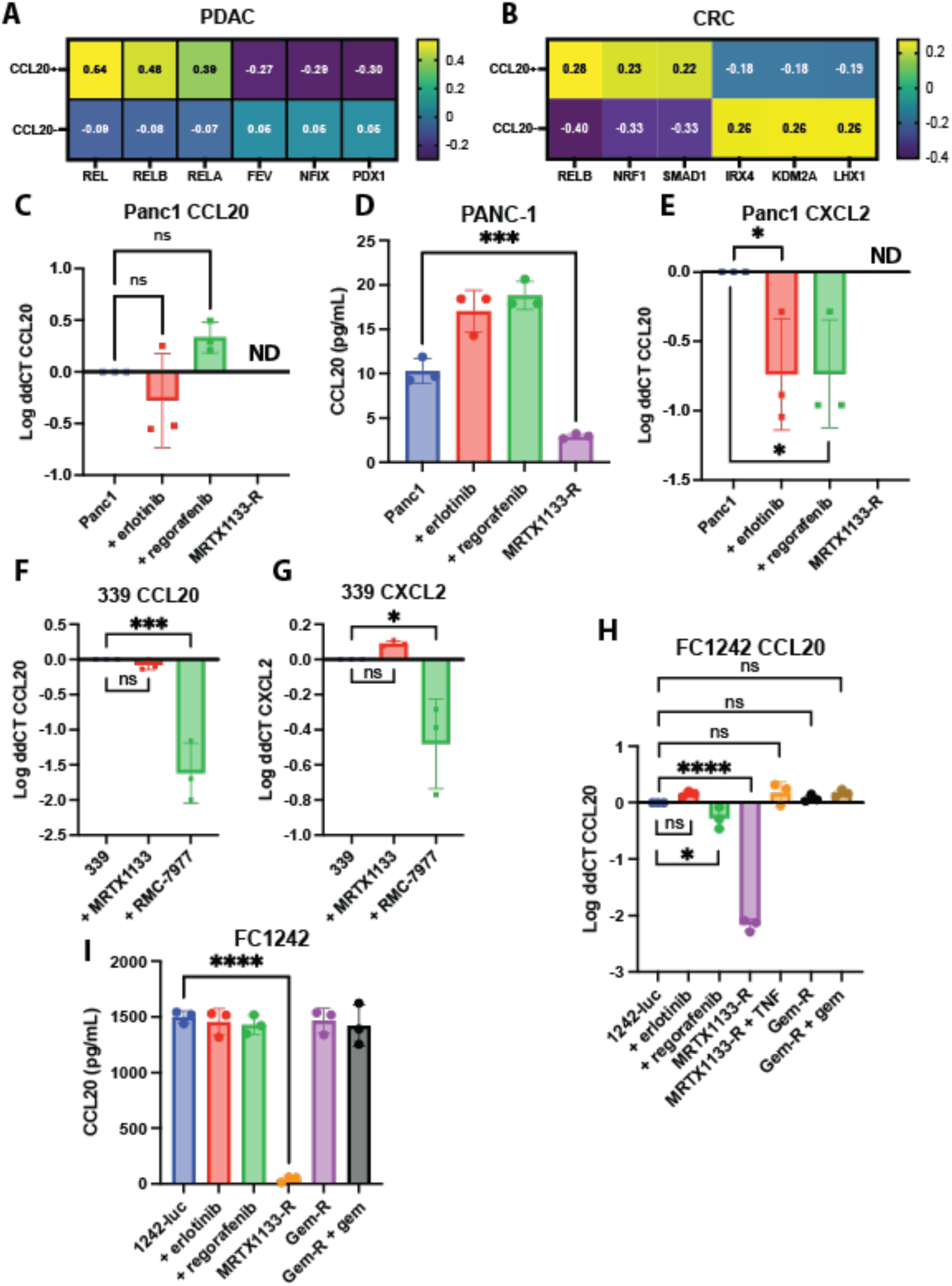
CCL20 expression is regulated by NF-κB and changes with KRAS inhibition. **A** The top three transcription factors inferred using CollecTRI^60^ from all pancreatic exocrine ductal cells in a single cell RNA dataset^61^ sorted by CCL20 expression. **B** Inferred transcription factors from tumor cells from a CRC dataset sorted by CCL20 expression. **C** qPCR and **D** ELISA for CCL20 transcript expression and protein secretion by human PANC-1 cell line. **E** qPCR for CXCL2 by the PANC-1 cell line. ND = not detected by qPCR. ns, not significant. **F** qPCR for CCL20 and **G** CXCL2 expression by the patient-derived 339 cell line possessing a KRAS^G12V^ mutation. **H** qPCR and **I** ELISA for CCL20 in the murine KPC FC1242 stably expressing Firefly luciferase PDAC cell line. Treatments were for 24 hours. Concentrations of the individual treatments were: Erlotinib 500 nM, regorafenib 500 nM, MRTX1133 500 nM, gemcitabine 100 nM, TNF 20 ng/mL with 10 ng/mL of IL-1, and RMC-7977 300 nM. For resistant cell lines (denoted by -R), resistance was considered established when cells survived in full growth medium supplemented with 10 μM of the denoted chemotherapy drug. One-way ANOVA was used for statistical analysis with Dunnett’s test for multiple comparisons used to compare to the untreated cell line.

### CCL20 transcript and protein changes in response to chemotherapies

Cytotoxic chemotherapies induce cell stress and cell death that activate NF-κB and result in downstream CCL20 expression in CRC and PDAC^23,66,67^. We set out to investigate if chemotherapy modulated CCL20 expression *in vitro*. For these studies we completed *in vitro* analysis of either human or mouse cell lines with key chemotherapies commonly used to treat patients with PDAC or CRC. The human PANC-1 cell line, possessing a single allele KRAS G12D mutation^68^, had low expression and protein detection of CCL20 at baseline (Fig. 4C, 4D), matching The Human Protein Atlas data (Fig. S3C)^69^. To test the pathway correlations that we found associated with CCL20 expression in Figure 3A and 3B, we first treated PANC-1 cells with erlotinib, an EGFR inhibitor approved in for use in PDAC^70^, or regorafenib, an RTK inhibitor that targets VEGFR1-3, TIE2, PDGFRB, FGFR, KIT, RET, and RAF and currently approved for use in CRC^71^. No changes in CCL20 were detected by ELISA or qPCR for PANC-1 cells treated with either erlotinib or regorafenib (Fig. 4C, 4D).

As KRAS inhibitors have emerged as powerful therapies in both PDAC and CRC, we next investigated if inhibiting KRAS modulated CCL20 expression. KRAS is the most common oncogene in PDAC, with almost 90% of PDAC patients possessing a KRAS mutation. Consistent with a role for oncogenic KRAS signaling in CCL20 transcript expression, both expression and protein production of CCL20 were significantly decreased in a PANC-1 cell line resistant to the KRAS G12D inhibitor MRTX1133^72^. Acute treatment could not be tested with these cells due to the large amounts of cell death we observed at low doses in treatment naïve cells. This apoptotic effect was not observed with erlotinib or regorafenib treatment *in vitro*. However, CXCL2 expression was lowered in PANC-1s treated with erlotinib or regorafenib or PANC-1s with MRTX1133 resistance (Fig. 4E). The patient-derived cell line MCW339, possessing a KRAS G12V mutation, had high basal level of CCL20 expression and protein that was lowered by acute treatment with the RAS Multi(ON) inhibitor RMC-7977^73^ but not the KRAS G12D inhibitor MRTX1133 (Fig. 4D, S4D, S4E). The same trend was observed for CXCL2 expression (Fig. 4G). These data suggest that mutant KRAS is the driving factor behind increased chemokine production in PDAC.

This reliance on oncogenic KRAS for CCL20 expression was further validated using murine PDAC cells harboring KRAS G12D mutations. While these cell lines had high baseline production of CCL20, expression of CXCL2 was undetectable (Fig. 4H, 4I). Gemcitabine resistance or the combination of gemcitabine resistance with gemcitabine added to the culture did not alter CCL20 expression or secretion. CCL20 expression lost in FC1242 MRTX1133-resistant cell line was restored following administration of TNF and IL-1. Similar patterns of basal and induced CCL20 expression were observed in the murine PDAC cell lines FC1199 and FC1245 (Fig. S4F-S4I). We next validated the CCL20 findings by analyzing a previously published scRNA sequencing dataset examining orthotopic KPC tumors^74^. From this dataset, we determined that expression of CCL20 and CXCL5, like CXCL2 is a CXCR1/2 ligand, was limited to the basal-like tumor cells with expression lost following KRAS G12D inhibition but not KRAS G12C inhibition (Fig. S4J). Lastly, there were no differences in CCL20 expression between different KRAS mutations in PDAC or CRC using the TCGA data (Fig, S4K, S4M).

### KRAS inhibitor resistance decreases phosphorylation of p65

Given the data suggesting NF-κB regulation of CCL20 expression in PDAC (Fig. 4A), we next sought to confirm that NF-κB signaling was downstream from oncogenic KRAS signaling. Using KRAS inhibitor treated or resistant cell lines, we examined phosphorylated p65 (RELA) by flow cytometry. As predicted, there was a significant decrease in phosphorylated p65 in either KRAS G12D-specific MRTX1133 resistant cells or the tri-complex RAS(ON) RMC-7977 resistant or treated (Fig. 5A, S5A, S5B). From these data we inferred that while mutant KRAS was not solely sufficient for elevated CCL20 and CXCL2 synthesis, mutant KRAS does appear to be necessary for NF-κB-mediated chemokine production in malignant pancreas epithelium. However, not all chemokines transcribed by NF-κB may be regulated in the same manner. For example, CXCL2 is transcribed by both NF-κB and FLI1 in an additive manner^64,75^ while FLI1 does not modulate CCL20 expression^76^. Thus, the reduction in NF-κB mediated transcription may be compensated for by FLI1 or other transcription factors independent of NF-κB, providing a mechanism for the differences we observed in CCL20 and CXCL2 expression.

**Figure 5.**
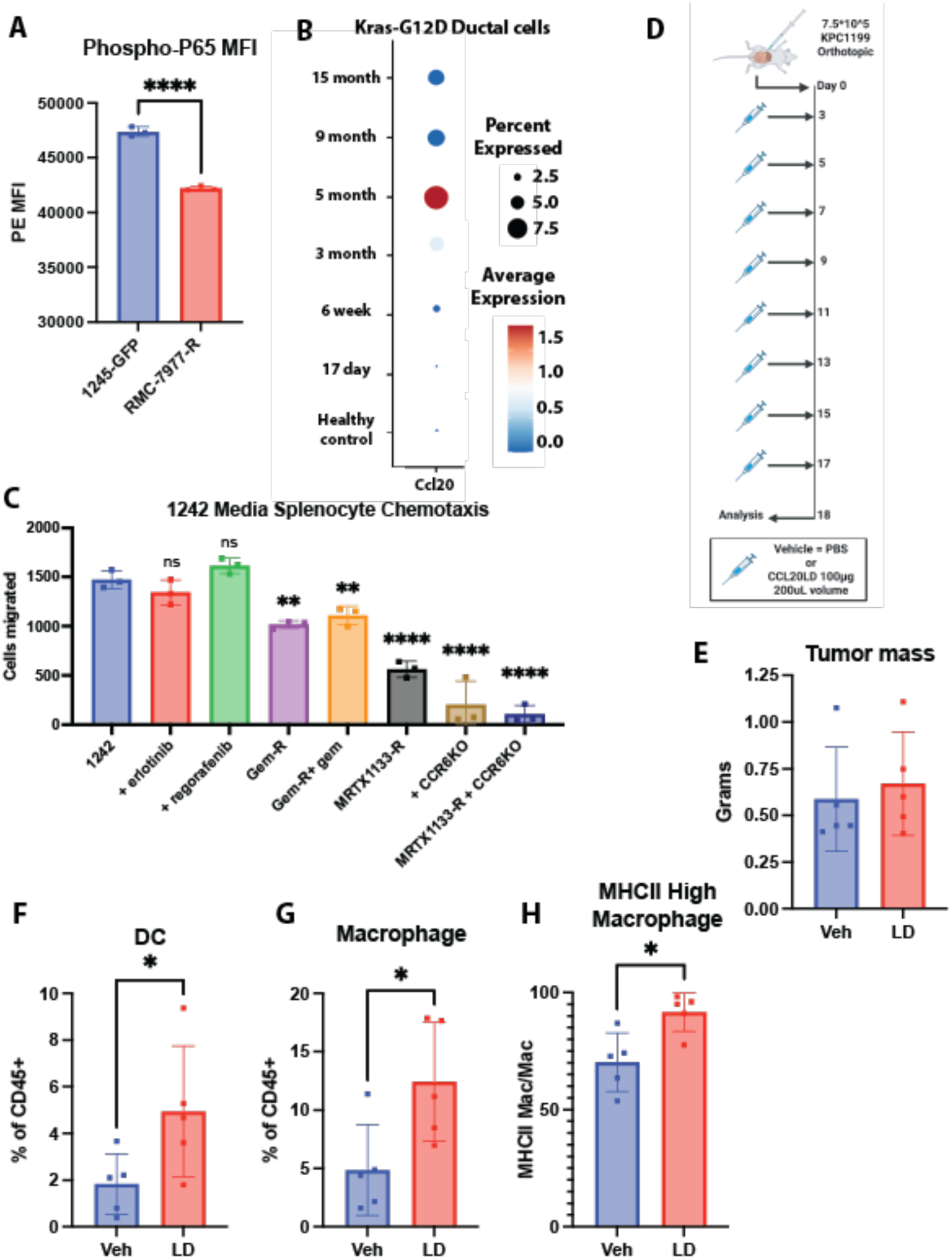
Oncogenic KRAS activates NF-κB and inhibition of CCR6 with CCL20LD increases infiltration of antigen presenting cells in PDAC. **A** Mean fluorescence intensity (MFI) of anti-phospho-p65-PE immunostained FC1245 and RMC-7977 (2 μM)-resistant FC1245 PDAC cells. * denotes *p* ≤ 0.05, Student’s unpaired *t*-test. **B** Expression of CCL20 in a single cell RNA dataset^78^ of pancreatic ductal cells obtained from mice with a *Ptf1a* specific tamoxifen inducible *Kras*-G12D. The time listed on the y-axis is time from when the initial tamoxifen was given. **C** The number of wild-type C57BL6 splenocytes migrated in a chemotaxis assay using conditioned media from the listed FC1242 PDAC cell line. The conditioned media was obtained from cells outlined in Figure 4I. One-way ANOVA was used for statistical analysis with Dunnett’s test for multiple comparisons used to compare to the untreated cell line **D** Treatment schema for mice treated implanted with orthotopic KPC1199 tumors and treated with either vehicle PBS or 100 μg CCL20LD. **E** The mass of the tumors from the experiment outlined in Figure 5D. **F** CD11c+ MHCII+ dendritic cells, **G** CD11b+ F4/80+ macrophages, and **H** MHCII high macrophages (as a percentage of macrophages) in the harvested tumors. * denotes *p* ≤ 0.05, Student’s unpaired *t*-test.

We have shown that loss of KRAS signaling caused loss of NF-κB driven CCL20 expression. Next, we wanted to investigate if activation of oncogenic KRAS elevates CCL20 expression. KRAS initially becomes mutated during PanIN progression and overactive G12D mutations accounting for around 40% of PDAC KRAS mutations^77^. To confirm if oncogenic KRAS could upregulate CCL20, we investigated a scRNA dataset taken from the pancreas of mice with a *Ptf1a*-specific tamoxifen-inducible mutant *KrasG12D*, a model of premalignant pancreatic intraepithelial neoplasia (PanIN)^78^. Here, we found CCL20 was, like PDAC, limited to the ductal cells (Fig. S5C). When assessing by time after tamoxifen injection, CCL20 expression increased from little to no transcript at 17 days or 6 weeks (Fig. 5B) to significant elevation CCL20 beginning at 3 months. This timeline is congruent with the known development of PanIN lesions in this mouse model^78^.

To test if the reduction in chemokine levels due to KRAS inhibitor resistance would result in decreased immune recruitment, we performed chemotaxis assays using conditioned media from the FC1242s and splenocytes from wild type C57BL/6J mice (Fig. 5C). Erlotinib and regorafenib treatment did not reduce the number of cells migrated. Gemcitabine resistance in the FC1242 cell line did reduce splenocyte migration, but this was not further lowered when gemcitabine was given to the gemcitabine-resistant cell lines 24 hours prior to media collection. The largest reduction in splenocyte chemotaxis was seen with MRTX1133 resistant cells. To see if CCL20 played a role in chemotactic migration reduction, we performed the same chemotaxis assay using splenocytes derived from CCR6 knock-out mice. In this assay, there were no differences in cell migration between wild type FC1242 and MRTX1133-resistant FC1242s, suggesting that CCL20 was a major factor accounting for the differences in splenocyte migration between the wild type and MRTX1133-resistant PDAC cell lines. Together, the data support the notion that CCL20 production was not altered by commonly used RTK inhibitors used to treat PDAC patients. However, the data revealed a dependency on mutant oncogenic KRAS signaling for CCL20 production. Oncogenic KRAS driven CCL20 expression was broadly observed across different KRAS variants, further supporting the notion that elevated CCL20 reflects over-activation of the KRAS monomeric G protein signaling pathway.

### Inhibition of CCR6 is not protective in orthotopic models of PDAC

To test if inhibition of oncogene-driven CCL20-CCR6 signaling would have a beneficial impact on modulating the PDAC TME, we used an orthotopic mouse model of PDAC treated with CCL20 locked dimer (CCL20LD), a chemokine variant we have shown to be CCR6 partial receptor agonist / inhibitor^18^. CCL20LD is a dimerized form of the wild type CCL20 that binds CCR6 with physiologic nanomolar affinity, but does not evoke chemotaxis, thus acting as a competitive orthosteric inhibitor of CCL20-CCR6 signaling. We have previously shown that CCL20LD has minimal side effects and a wide therapeutic window in healthy mice, making it a good candidate for translation to the clinic as a precision immunotherapy^19^. Tumors were implanted and mice were treated with either vehicle PBS or 100 μg CCL20LD over the 18 days prior to tumor harvest (Fig. 5D). While there were no significant differences in tumor size or volume between the two groups (Fig. 5E), there were significant increases in CD11c+ MHC class II high dendritic cells, F4/80+ macrophages, and MHCII high macrophages in the CCL20LD treated group (Fig. 5F-5H). Further analysis of the TME by flow cytometry did not detect a difference in the relative frequencies of leukocytes, CD4+ T cells, CD8+ T cells, CD11b+ myeloid cells, CD19+ B cells, and CD3+ T cells (Fig. S5D-S5I). Surprisingly given their robust expression of CCR6, levels of tumor infiltrating IL-17A+ Th17s and FOXP3+ Tregs were unchanged (Fig. S5J, S5K). Lastly, the two major subtypes of Tregs (RORγT+ and Helios+) were unchanged as well (Fig. S5L, S5M). While our data match our previous report showing increased circulating macrophages and dendritic cells in healthy mice administered with comparable doses of CCL20LD^19^, they are at odds with previous reports of decreased macrophages in colorectal cancer tumor engrafted to CCR6 knockout mice^79^. The discrepancy between these results may reflect acute compared to lifelong loss of CCR6 function as CCR6 knockout mice have dysregulated localization of dendritic cells and underdeveloped gastrointestinal lymph nodes from birth^80^. Ultimately, these findings indicate that acute therapeutic inhibition of CCL20-CCR6 signaling in established PDAC tumors increased the infiltration of antigen presenting DCs and macrophages within PDAC tumors.

## DISCUSSION

Attempts to target the immunosuppressive TME in PDAC have not yielded many successful strategies^81^. Targeting chemokine signaling has shown promise but the lack of therapeutics to target chemokine ligands and receptors outside of CXCR1 and CXCR2 remain little explored. To identify other, non-CXCR1/2 therapeutic targets, we screened for chemokine ligands upregulated in PDAC. We find that CCL20 is upregulated in PanINs and human PDAC, suggesting it may contribute to the initial or early formation and maintenance of the immunosuppressive TME. Through extensive computational analysis with confirmation *in vitro*, we show that the CCL20 is driven by NF-κB signaling. We identified this signaling was limited to the basal subtype and found that CCL20 expression was lost with resistance to KRAS inhibitors. Once cells are resistant to KRAS inhibitors, their chemokine expression decreased resulting in decreased leukocyte recruitment. Lastly, through use of a novel orthosteric inhibitor of CCR6 we demonstrated that inhibiting CCL20-CCR6 signaling increased tumor infiltration of antigen-presenting cells, as summarized in Figure 6. Cumulatively, these data illustrate tumor specific and cancer subtype specific mechanisms influencing CCL20 expression and outline a precision immunotherapy strategy for targeting CCL20 in PDAC.

**Figure 6.**
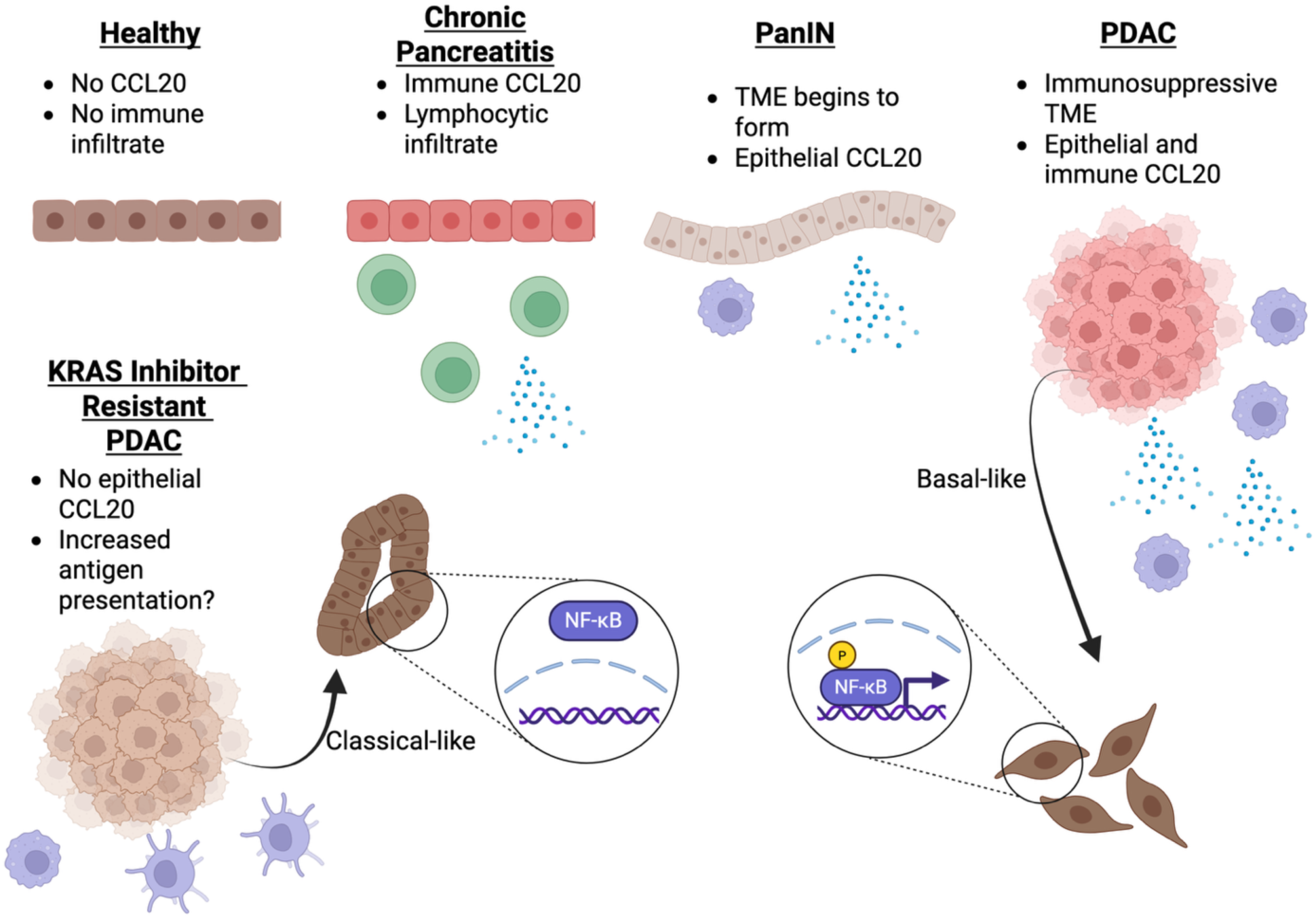
The evolution of CCL20 production in the diseased pancreas. **A** Graphical abstract summarizing the findings. CCL20 is not expressed in the healthy pancreas. In pancreatitis, it is expressed the lymphocytic infiltrate. In PanINs, CCL20 begins to be expressed by the ductal epithelium. In PDAC, CCL20 expression is further increased by the ductal tumor cells and infiltrating myeloid cells. CCL20 expression is lost with resistance to KRAS inhibitors, along with decreased phosphorylation of *RELA*, and is limited to the basal subtype. Inhibition of CCL20-CCR6 signaling increased levels of antigen presenting cells, including dendritic cells and MHCII-high macrophages within the tumor parenchyma.

CCL20 is known to be regulated by NF-κB which we confirmed *in silico*. Our data indicate that loss of CCL20 with development of resistance or treatment with KRAS inhibitors suggests that mutant KRAS is activating NF-κB signaling and in turn driving CCL20 production. KRAS activating NF-κB has been reported before although the mechanisms proposed have been varied. Hypothesized signaling pathways include KRAS-mediated overexpression of GSK3a upstream of IKK^82^, release of IL-1 causing autocrine NF-κB activation^83^, or activation of PI3K resulting in AKT and NF-κB activation^84^. In our study, we did not determine the specific mechanisms in which KRAS activates NF-κB. However, we did find oncogenic KRAS-dependent activation of canonical NF-κB signaling. Constitutive canonical NF-κB signaling has been found in PDAC previously^85^. Interestingly, our transcription factor analyses implicated both canonical and non-canonical NF-κB signaling driving CCL20 expression. A recent study found that non-canonical RELB is only able to bind to open chromatin while canonical RELA is able bind both open and closed chromatin^86^. Our epigenetic analysis of CCL20 suggested that it is likely in an open chromatin region in PDAC, making it possible to be transcribed by both canonical and non-canonical NF-κB signaling. Further work is necessary to identify if oncogenic KRAS can activate both canonical and non-canonical NF-κB signaling in PDAC.

In a study by Li et al., CCL20 and other NF-κB chemokines produced by tumor cells were linked to T cell low clones in mouse models of PDAC^87^. However, the authors attributed the increased chemokine expression to MYC overexpression and chromatin accessibility but did not investigate a link to NF-κB. MYC amplification has been found to be a common mechanism of resistance to RAS inhibition, but we did not observe a concurrent upregulation of CCL20 or CXCL2 in the RAS inhibitor resistant cell lines^88^. Although resistance to KRAS inhibitors develops for most patients^89^, our data suggest there is a large benefit to reducing KRAS and thus NF-κB signaling in cancer cells. The loss of NF-κB driven chemokines can promote a more favorable TME that may be conducive to combination treatment with immunotherapies^90,91^. Lastly, NF-κB plays a large role in traditional cytotoxic chemoresistance which supports preclinical data showing improved tumor control with combinatorial KRAS inhibition and chemotherapy^92^. Interestingly, mutant KRAS is associated with the iCMS3 subtype of CRC, not the iCMS2 subtype of CRC that we found correlated with CCL20 expression^48^. The iCMS2 subtype is enriched for *APC* and *TP53* mutations, which may indicate that loss of function with P53 may drive CCL20 expression, the cell of origin and its epigenetic profile may play a role in regulating expression, external cues from the microbiome can stimulate CCL20 expression, or that multiple oncogenes can promote CCL20 expression through different pathways that likely converge on NF-κB.

CCL20 has been linked to the classical subtype of PDAC, but this link was established using 10X Visium data which lacks single cell resolution^93^. Further, our basal subtype specific expression of CCL20 that is lost with KRAS inhibition is in line with current data that show KRAS inhibitors preferentially target the basal and not classical subtype of PDAC^92^. A recent study suggested that the transcription factor specific protein 5 (SP5) regulates CCL20 expression in PDAC^94^. However, while SP5 was not observed in our transcription factor analyses, SP5 is also known to repress the classical subtype transcription factor GATA6^95,96^. Further analyses of SP5 in a non-PDAC cells lacking GATA6 repression, such as macrophages, will benefit the results as it may help to further tease apart the distinction between PDAC subtypes influencing CCL20 expression and necessary transcription factors for CCL20. CCL20’s sole receptor is CCR6, whose expression is predominant on Th17 cells, Tregs, and immature DCs. Like CCL20, CCR6 has been variably described as tumor promoting or inhibiting^97^. As Th17 cells have been linked with PDAC initiation or promotion or progression, we predicted that the locked dimer variant, CCL20LD would result in smaller tumors *in vivo*. Supporting this postulate, prior work has shown that treatment with CCL20LD potently inhibits the trafficking of CCR6+ T cells into psoriasis or psoriatic arthritis lesions^6,18^. Surprisingly, CCL20LD treatment of orthotopically engrafted PDAC resulted in increased dendritic cells within the tumor when used as a monotherapy, independent of reduction in tumor size or changes in T cell infiltration. Given that dendritic cells present tumor antigens to cytotoxic lymphocytes and are known to sensitize tumors to checkpoint blockade it remains possible that KRAS-dependent CCL20 production influences immune evasion by excluding those cells from the tumor parenchyma^98^. Thus, future studies examining CCL20LD should do so as a combinatorial therapy with other immunomodulators such as anti-PD-1 immune checkpoint blockade. Another possibility would be to perform further subtyping of the dendritic cells present to identify if the infiltrating dendritic cells were anti-tumor cDC1s or to perform immunopeptidomics to see if CCL20LD increased presentation of tumor-specific antigens^99^. Alternatively, dendritic cells have shown to be protective in pancreatitis^98^ and CCR6+ Th17 cells are known to play a role in chronic pancreatitis pathogenesis^100^. Thus, there may be an opportunity for CCL20LD to be used as a preventative or therapeutic for chronic pancreatitis, one of the largest risk factors for developing PDAC^101^.

In summary, our data thoroughly characterized the cell types that express CCL20 during a healthy, inflamed, pre-malignant, and malignant state in the pancreas. We defined when CCL20 is upregulated and downregulated by identifying specific subtypes and therapies that changed CCL20 production. We identified the impact of inhibiting CCL20-CCR6 signaling in the PDAC TME using CCL20LD, a compound we have previously used as an immunomodulatory therapeutic for inflammatory diseases such as psoriasis and psoriatic arthritis^6,8,18^. The increases in antigen presentation seen with CCL20LD suggest that its immunomodulatory benefits could extend from being an anti-inflammatory for use in autoimmune diseases to also providing benefit as a precision immunotherapy for cancer. Altogether, our data identify a potential window for precision chemokine-targeted immunotherapy that selectively disrupts aberrant chemokine production downstream of oncogenic KRAS in PDAC.

## METHODS

### Mining single cell RNA and bulk sequencing datasets

The list of single cell RNA datasets used, their GEO accession number, and corresponding papers can be seen in Supplementary Data 6. The single cell datasets were analyzed using R 4.4.3 (RRID:SCR_001905), RStudio (RRID:SCR_000432) 2024.12.1+563, and Seurat 5.2.1 (RRID:SCR_016341)^102^. Differential gene expression was performed using MAST^103^ with a minimum percent expression cut-off of 5% and a log fold change threshold of 0.2. The differentially expressed genes based on disease state can be seen in Supplementary Data 3. The analysis of TCGA and GTEx bulk sequencing data was performed with GEPIA^104^. Transcription factor analysis was performed on single cell datasets using CollecTRI package^60^. General transcription factor analysis was performed using ContraV3^63^ analyzing 500 base pairs upstream of the promoter. The gene sets used for the correlation analyses in the single cell RNA datasets can be seen in Supplementary Data 7. Methylation of the CCL20 gene was performed using cBioPortal^59^ using the CCL20 cg12643045 probe.

### Cell culture

Human HPNE (RRID:CVCL_C466), Capan-2 (RRID:CVCL_0026), MIA PaCa-2 (RRID:CVCL_0428), PANC-1 (RRID:CVCL_0480), HCT116 (RRID:CVCL_0291), and AsPC-1 (RRID:CVCL_0152) cells were obtained from the ATCC (Manassas, VA). The human pancreatic stellate cell line (HPSC) was derived from human PDAC tumor and immortalized with hTERT and SV40T plasmids under Neomycin antibiotic selection and used to model inflammatory cancer-associated fibroblasts as defined previously^105^. The patient-derived MCW339 and MCW512 cells obtained from consenting patients receiving surgical resection as part of their standard of care in the LaBahn Pancreatic Cancer Program and were de-identified of patient identifiers and provided by the Medical College of Wisconsin Ronald Burklund Eich Biorepository in accordance with an IRB approved protocol. As detailed in our prior publications^40,106^, patient-derived cells were isolated following manual and enzymatic digestion from consented patient specimens of primary pancreatic ductal adenocarcinomas, serially diluted and cultured in 2D tissue culture dishes in DMEM-F12 media (ThermoFisher, Carlsbad, CA #11320033) containing 10% (v/v) fetal bovine serum (FBS) (Omega Scientific, Tarzana, CA #FB-12), penicillin/streptomycin (Pen/Strep) (ThermoFisher #15140122) 100 U insulin (Millipore Sigma, Milwaukee, WI, #I6634), bovine pituitary extract (ThermoFisher #13028-014), epidermal growth factor (ThermoFisher #PHG0313) and hydrocortisone (Millipore Sigma #H0888), and L-Glut (ThermoFisher #25030-081).

The murine pancreatic cancer cell lines FC1199, FC1242, FC1245, and DT10022 were derived from founder KRas^LSL.G12D/+^-p53^R172H/+^-Pdx^Cre^ (KPC) mice on a homogenous C57BL/6 background and were kindly provided by Dr. Dannielle Engle and Dr. David Tuveson (Cold Spring Harbor Laboratories, NY). The HCT116 DNMT knockouts were previously provided by the laboratory of Dr. Bert Vogelstein^107^ (John Hopkins University School of Medicine, MD). The HPSC, MIA PaCa-2, PANC-1, and all murine cell lines were cultured in DMEM (ThermoFisher #11965-092) supplemented with 10% (v/v) FBS. AsPC-1 cells were cultured with RPMI-1640 (ATCC #30-2001) supplemented with 10% (v/v) FBS. Capan-2 and HCT116 cells were cultured with McCoy’s 5A medium (ThermoFisher #16600-082) supplemented with 10% (v/v) FBS. The HPNE cells were cultured with DMEM (Millipore Sigma #D-5030) dissolved into 1 L ddH_2_O supplemented with 10 mL L-Glutamine, and 20 mL sodium carbonate (ThermoFisher #BP357-1). The DMEM solution was then filter sterilized and 375 mL of sterile DMEM solution was supplemented with 125 mL M3:BaseF (INCELL, San Antonio, TX #M300-F), 75 mL FBS, 5 mL Pen/Strep, 2.75 mL of 1 M D-Glucose (ThermoFisher #J60067-AK), 50 μL epidermal growth factor, and 37.5 μL puromycin (ThermoFisher #A1113803). Cell lines were authenticated annually using short tandem repeat profiling and mycoplasma-tested semi-annually.

### Creation of drug-resistant cell lines

Drug-resistant cell lines were created by initial dosing of the cell lines with 1 nM or below of the specified drug. Cells were treated four days with each inhibitor, followed by passage of the surviving cells, recovery growth in full medium without the inhibitor for three days, and then application of double the previous concentration of drug for additional four days. This process was repeated until cells were able to grow in the presence of 10 μM of the specified drug, except for the RAS(ON) inhibitor RMC-7977 (Selleckchem, Houston, TX, #E1858) which was stopped at 2 μM. Gemcitabine (Hospira, Pleasant Prairie, WI) and MRTX1133 (MedChemExpress, Monmouth Junction, NJ, #HY-134813) were used.

### ELISAs

Cells were cultured in a 6-well tissue culture plate (Corning, Corning, NY, #3506). Upon reaching 80% confluence, the medium was changed and the specified drug was added. Cells were then allowed to grow for 24 hours prior to media collection. Drug concentrations included 500 nM MRTX1133, 500 nM erlotininb (MedChemExpress #HY-50896), 500 nM regorafenib (MedChemExpress HY-10331), 100 nM gemcitabine, or 20 ng/mL TNF (ThermoFisher #300-01A) and 10 ng/mL IL-1 (ThermoFisher #200-01A). The ELISAs were performed using the human CCL20 ELISA (R&D Systems, Minneapolis, MN, #DY360) or mouse CCL20 ELISA (R&D Systems #DY760) following the manufacturer’s protocol. Absorbance was measured on BioTek Cytation 5 Cell Imaging Multi-Mode Reader (RRID:SCR_019732). Extrapolation of the values was performed by using non-linear or linear regression using GraphPad Prism (RRID:SCR_002798). All R^2^ values of the standard curves were > 0.96.

### PCR and qPCR

The list of primers and cycling conditions used for PCR and qPCR can be seen in Supplementary Data 8. RNA was obtained from cells using the RNeasy Mini Kit (Qiagen, Hilden, Germany, #74104) following the product protocol. RNA was collected at the same point as the conditioned media for the ELISAs. RNA was converted to cDNA using a reverse transcription kit (ThermoFisher #4368814). qPCR was performed using the SsoAdvanced supermix (BioRad, Des Plaines, IL, #172-5282). 250nM of the forward and reverse primers were used along with 150nM of the fluorogenic probe and 1 μL of cDNA in a 10 μL reaction for qPCR. The reaction was performed using the CFX Opus 96 (BioRad) and analyzed on CFX Manager (RRID:SCR_017251).

### Immunofluorescence

Five PDAC tumor formalin fixed paraffin embedded (FFPE) sections, and five healthy pancreas sections were obtained from different patients undergoing surgical resection as part of their standard-of-care and obtained from the Medical College of Wisconsin Ronald Burklund Eich Biorepository. Samples were deparaffinized, blocked with 2% (v/v) goat serum (ThermoFisher #31872), stained with mouse anti-human CCL20 (R&D systems MAB360) at a concentration of 1:100 and goat anti-mouse IgG Alexa fluor 647 (ThermoFisher #A48289) at a concentration of 1:500. A mouse IgG1 (R&D Systems #MAB002) was used as an isotype control. Sections were counterstained with a mountant containing Hoechst 33342 (ThermoFisher #P36983). Slides were imaged on a Zeiss LSM 510 Confocal Microscope (RRID:SCR_018062). CCL20 was quantified using ImageJ (RRID:SCR_003070). The percent of CCL20 immunostained cells within the Hoechst area was used to account for non-specific staining of red blood cells within at least three fields of view per sample. The average CCL20+ area per sample was calculated and graphed.

### Mutation and copy number signatures

Mutation annotation files (MAF) were obtained from the Genomics Data Commons data portal. Single base substitution (SBS) signatures were obtained from the MAF files using SigProfilerMatrixGenerator^108^ in Conda (RRID:SCR_018317). All SBS signatures that were detected in >10% of PDAC or CRC samples were then used to divide TCGA samples into groups and analyzed by unpaired *t*-test using the CCL20 z-score (RSEM) obtained from cBioPortal. All SBS signatures from the TCGA patients can be seen in Supplementary Data 5 (PDAC) and Supplementary Data 6 (CRC). Copy number signatures were obtained from the work of Steele et al.^58^ and analyzed with similar methods as the SBS signatures. The SBS and CN signature groups created from cBioPortal can be accessed through the link in Supplementary Data 9.

### Mouse use

All animal experiments were conducted under approved protocols from the Medical College of Wisconsin (AUA00076) and in accordance with the National Institutes of Health Guide for the Care and Use of Laboratory Animals and the ARRIVE guidelines^109^. Male C57BL/6 mice (RRID:IMSR_JAX:000664) were purchased from the Jackson Laboratory (Bar Harbor, ME). Animals were maintained on a strict 12:12 hour light–dark cycle in a temperature and humidity controlled facility with water and food provided *ad libitum*. Mice were monitored twice per day and weighed prior to the start of the experiment and then once weekly during the experiment. If an animal was determined to be in overt pain/distress beyond the point where recovery seemed reasonable, the animal was humanely euthanized in accordance with the *American Veterinary Medical Association Guidelines on Euthanasia*.

### Recombinant engineered chemokine

CCL20LD was provided by XLock Biosciences, Inc., Milwaukee, WI and was expressed and purified as previously described^18^. Protein purity was determined by mass spectrometry, and biologic function was confirmed by the inhibition of wild-type CCL20–dependent migration of CCR6-transfected Jurkat cells (RRID:CVCL_U617) in a Transwell chemotaxis assay prior to *in vivo* experiments. A concentration of 100 μg per mouse, previously shown to be effective in treating mouse models of psoriasis while exhibiting minimal side effects, was chosen for the treatment of tumor-bearing mice^6,18,19^.

### Orthotopic PDAC tumors

Six-to-eight-week-old male C57BL/6 mice were randomly assigned to treatment or control groups. Under anesthesia, 75,000 FC1199 cells were implanted into the head of the pancreas in 30 μL of DMEM media (day 0). Mice received injections of 100 μg CCL20LD in 200 μL PBS or 200 μL PBS as a control. Injections were every other day from days 3-17 for a total of 7 injections. Tumors were excised and weighed post-mortem.

### Flow cytometry

Tumors were digested following harvest using the mouse tumor dissociation kit (Miltenyi Biotec, Bergisch Gladbach, Germany, #130-096-730). For internal staining, cells were permeabilized with the FOXP3 transcription factor staining kit (ThermoFisher #00-5523-00). The antibody panel used for staining can be seen in Supplementary Data 10. Samples were run on the Cytek Aurora Spectral Analyzer (RRID:SCR_019826) and analyzed with FlowJo (RRID:SCR_008520). For the phosphorylated p65 flow cytometry, cells were serum starved in 0.5% (v/v) FBS media 12 hours prior to cell harvest and internal immunostaining.

### Chemotaxis assays

Conditioned media was gathered from PDAC cell lines as described previously. 235 μL of conditioned media was put into the bottom of 96 well Transwell plates (Corning #3387). Spleens were excised from euthanized mice, minced, and treated with a red blood cell lysis buffer (Biolegend #420301). The splenocytes were washed and filtered through a 70 μm strainer (Greiner, Kremsmunster, Austria, #542070). 100,000 splenocytes in 80 μL of serum-free RPMI with 0.2% (w/v) BSA fraction V (Thermofisher #15260-037) were loaded into the top of the Transwell plates. The plate was incubated at 37C for three hours and the number of cells in the bottom wells were analyzed using the Novocyte Advanteon (RRID:SCR_019522). CCR6 knock-out splenocytes were obtained from CCR6 knock-out mice (RRID:IMSR_JAX:005793).

## ACKNOWLEDGEMENTS

We thank Galina Petrova, PhD for her guidance and training in the Medical College of Wisconsin’s Children’s Research Institute Flow Cytometry Shared Resource. This research was completed in part with computational resources and technical support provided by the Research Computing Center at the Medical College of Wisconsin.

## DECLARATIONS

### Author Contributions

D.D. and M.B.D. wrote the main manuscript text. D.D. prepared each of the figures. D.D., M.D., and D.M. completed experiments and analyses. All authors reviewed the manuscript.

### Data Availability Statement

All data relevant to the study are included in the article. Any further information about resources and reagents should be directly requested to the corresponding author and will be fulfilled on reasonable request.

### Ethical Approval

Preclinical experimental procedures involving animals were conducted in accordance with the NIH Guide for the Care and Use of Laboratory Animals and were formally reviewed and approved by the Medical College of Wisconsin Institutional Animal Care and Use Committee (IACUC) (approval No. AUA000076). Use of de-identified tissue samples were obtained in accordance with an exempt IRB protocol (approval No. 234-03.

