## Supplementary material for "Mutant KRAS promotes NF-κB driven CCL20 chemokine expression in pancreatic ductal adenocarcinoma": Supplmental Figures

### Supplemental Figure Legends

#### Figure S1. CCL20 predicts a poor prognosis in PDAC but a favorable prognosis in CRC.

**A** Kaplan-Meier curve from TCGA PDAC patients or **B** CRC patients sorted by CCL20 expression.

#### Figure S2. Mutation signature analysis of PDAC and CRC patients.

**A** Expression of APEX1 and LIG1 in PDAC (left) and CRC (right) patients and sorted by SBS1 score of patient samples within the TCGA database. **B** KRAS mutation frequency (far left) and diagnosis age (middle left) in PDAC patients sorted by SBS1 and SBS5 (right) score from TCGA data. **C** KRAS mutation frequency in CRC patients by SBS1 (left) and SBS5 (right) score from TCGA data. Panel A used Student's unpaired *t*-test for statistical analysis. Panels B and C used Wilcoxon test for statistical analysis.

#### Figure S3. DNMT status with CCL20 expression.

**A** qPCR and **B** ELISA for CCL20 in the HCT116 cell line. **C** CCL20 normalized transcripts per million from the Human Protein Atlas for various CRC and PDAC cell lines. **D** qPCR for CXCL2 in the HCT116 cell line. One-way ANOVA with Dunnet's multiple comparison test to the wildtype used for S3A, S3B, and S3D. * denotes $p \leq 0.05$.

#### Figure S4. NF-κB and KRAS signaling increase CCL20 expression.

**A** ContraV3 results from transcription factors regulating CCL20 expression. **B** CXCL2, CXCL3, CXCL5, and CXCL8 expression from tumor cells sorted by CCL20 expression from a scRNA atlas of PDAC. **C** CXCL2 and CXCL3 expression of tumor cells from a CRC dataset sorted by CCL20 expression. All displayed chemokines were significantly increased in the CCL20+ tumor group by MAST analysis. **D** ELISA for CCL20 in the patient-derived 339 cell line, possessing a G12V mutation, receiving MRTX1133 or **E** RMC-7977. **F** ELISA and **G** qPCR for CCL20 in the mouse KPC FC1199 cell line. **H** qPCR and **I** ELISA for CCL20 in the mouse KPC FC1245-GFP cell line and a 2 μM RMC-7977 resistant FC1245-GFP cell line. **J** Expression of CCL20 and CXCL5 from a KPC (KRAS G12D) PDAC orthotopic tumor scRNA dataset treated with either vehicle, a KRAS G12C inhibitor, or a KRAS G12D inhibitor. **K** Normalized CCL20 expression, sorted by type of KRAS mutation, in TCGA PDAC and **L** TCGA CRC data. Panels S4D,

E, H, and I use student’s unpaired t-test. Panels S4F and S4G use one-way ANOVA with Dunnet’s multiple comparison test to the untreated cell line. * denotes $p \leq 0.05$.

**Figure S5. CCL20LD does not change lymphocyte populations in the PDAC TME.**

**A** Phospho-p65 MFI in PANC-1 and **B** MCW339 cells. **C** Expression of CCL20 in a single cell RNA dataset of pancreatic cells obtained from mice with a *Ptf1a* specific tamoxifen inducible *Kras*-G12D. **D** Mice were treated with 100ug of CCL20LD or vehicle 7-times over a span of 18 days following orthotopic tumor implantation with the FC1199 cell line. The tumors were harvested on day 18 and processed for flow cytometry. **D** CD45+ cells, **E** CD4+ T cells, **F** CD8+ T cells, **G** CD11b+ myeloid cells, **H** CD3+ T cells, **I** CD19+ B cells, **J** CD4+ IL-17A+ Th17 cells, and **K** CD4+ FOXP3+ Treg cells as a percentage of CD45+ cells. **L** ROR$\gamma$T+ and **M** Helios+ Tregs as a percentage of total Tregs. One-way ANOVA with Dunnet’s multiple comparison test to the wildtype used for S5A. Student’s unpaired *t*-test used for S6B, S6D-S6H. * denotes $p \leq 0.05$. No significant differences were observed by Student’s unpaired *t*-test in figures S6D-S6H.

**Supplemental Figure 1**

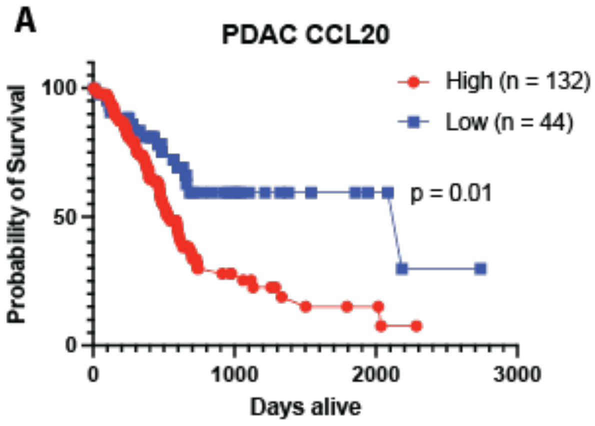


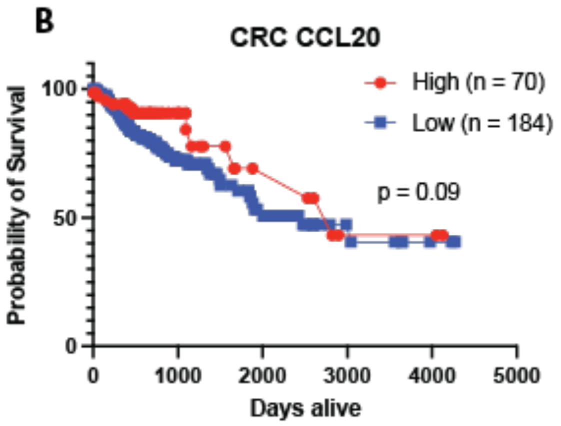

**Supplemental Figure 2.**

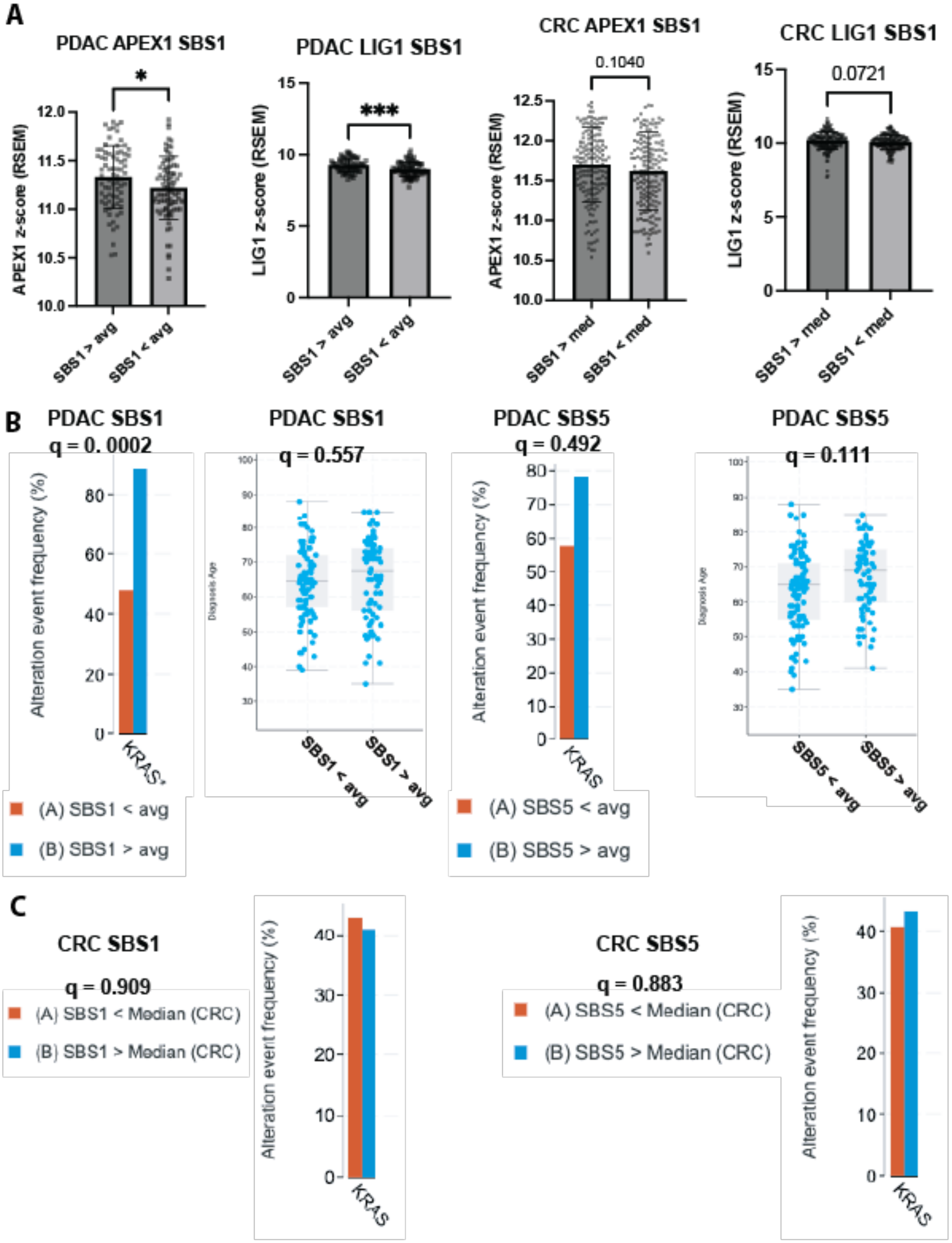

**Supplemental Figure 3.**

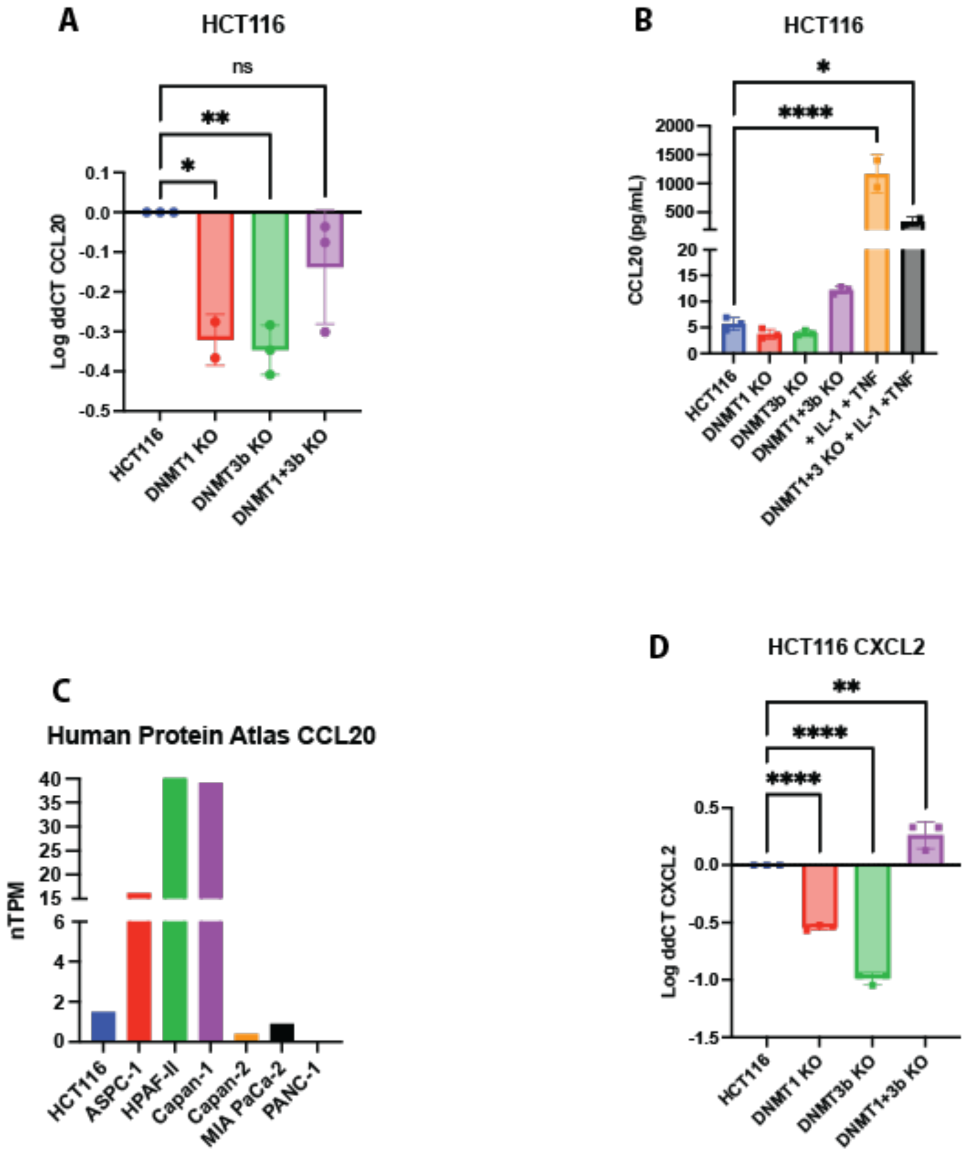

### Supplement Figure 4

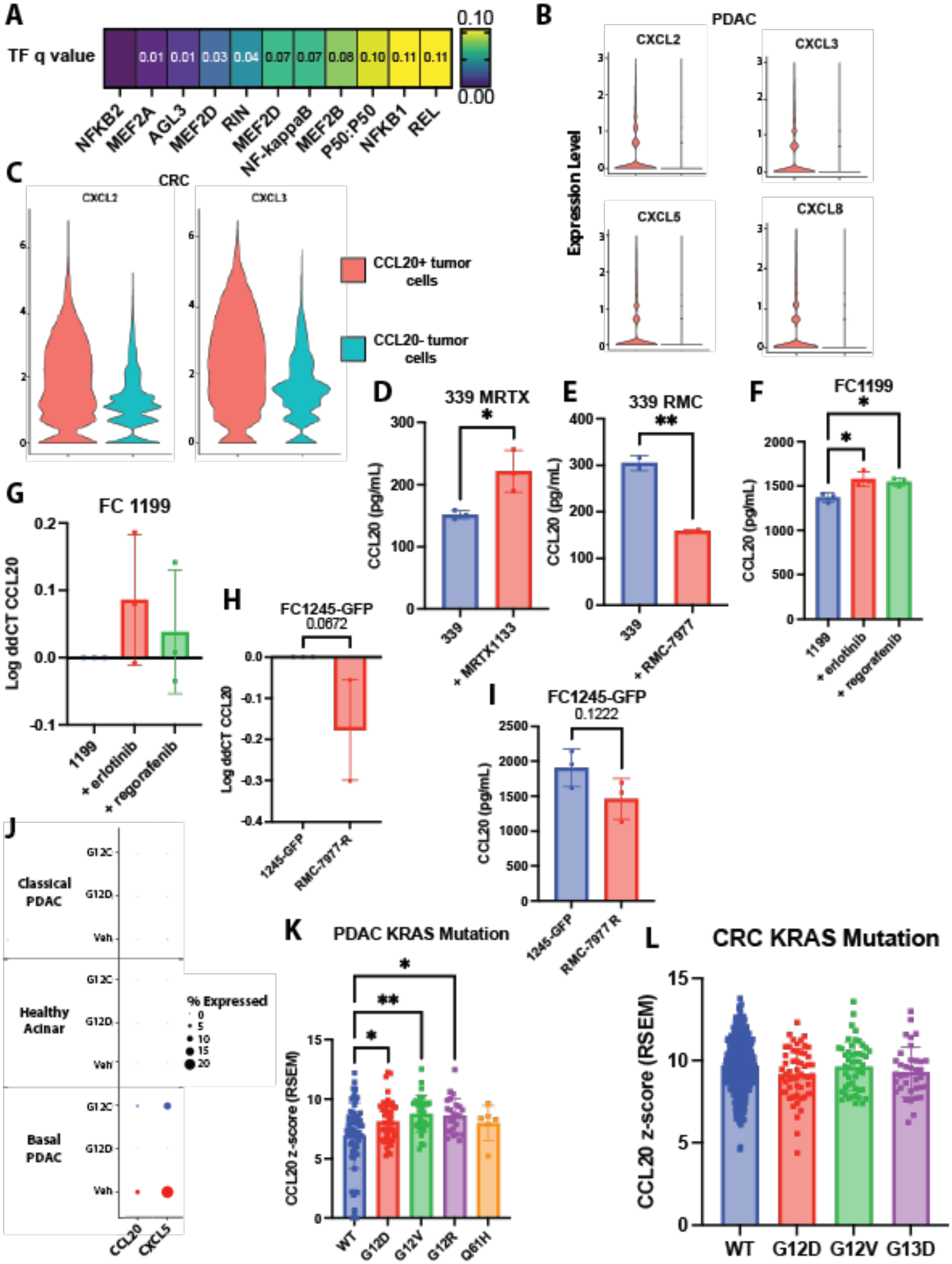

### Supplemental Figure 5

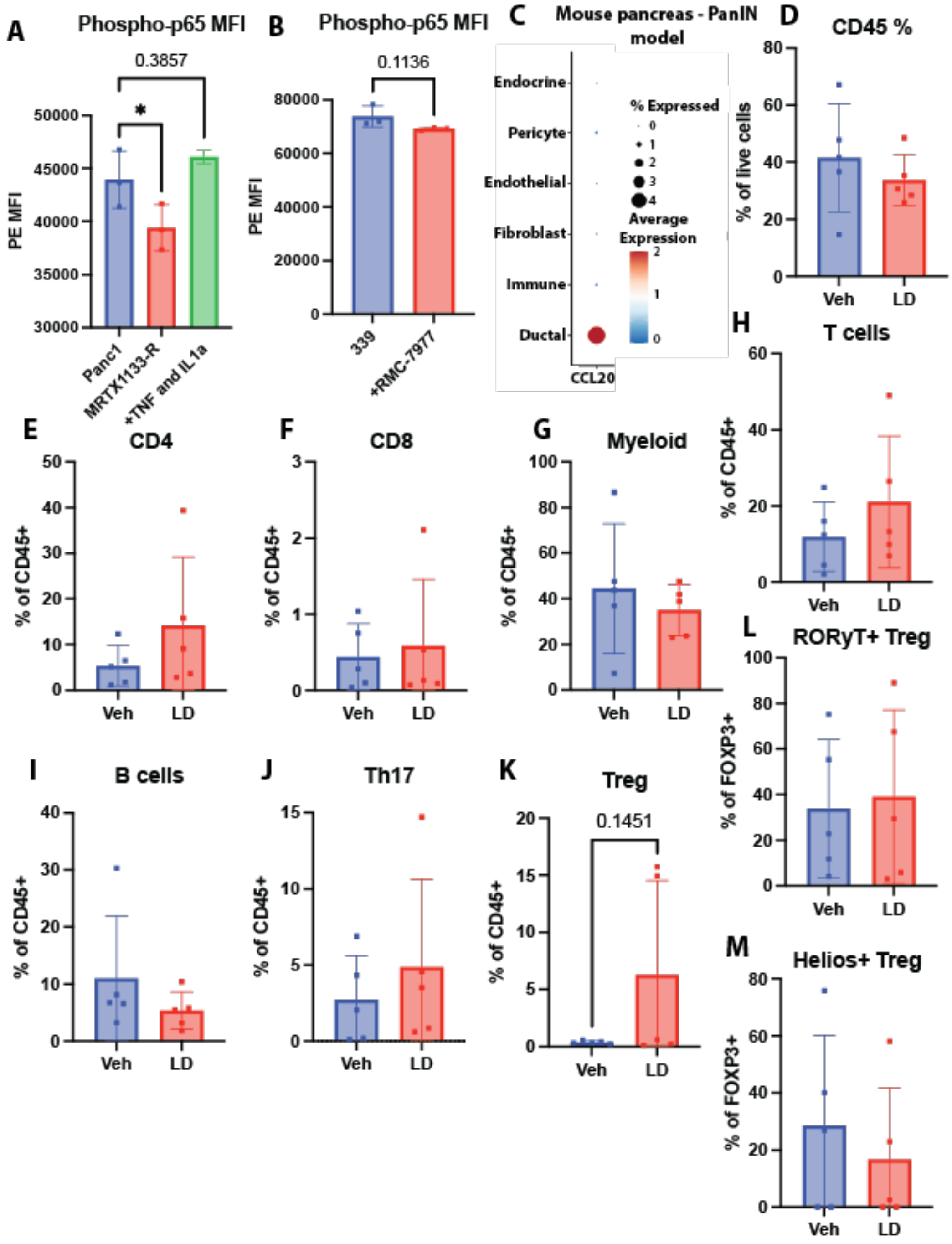
